# Contractile forces direct the chiral swirling of minimal cell collectives

**DOI:** 10.1101/2024.09.02.610752

**Authors:** Ghina Badih, Alexandre Schaeffer, Benoît Vianay, Pauline Smilovici, Laurent Blanchoin, Manuel Thery, Laëtitia Kurzawa

## Abstract

Chirality is a conserved biological feature with critical implications in tissue morphogenesis and embryonic development. In culture, large multicellular groups exhibit spontaneous chiral symmetry break when moving collectively on micropatterned surfaces. Although several studies have shown that actin network integrity and acto-myosin network contractility participate to the establishment of the chirality of the movement, the exact contribution of contractile forces to the directionality of the chiral bias in collectives remains to be elucidated. Here we studied the contractile forces produced by a minimal collective constituted of a pair of endothelial cells. We first show that cell doublets confined on disk-shaped micropatterns undergo spontaneous and persistent chiral swirling, displaying a mild but robust clockwise (CW) bias, as the one observed in bigger collectives. This bias could be amplified or reversed by modulating contractile forces. Traction force measurements revealed that large forces tend to drive counter-clockwise (CCW) rotation whereas low forces rather favor a CW rotation. Furthermore, the study of heterotypic doublets indicates that the speed and direction of the rotation is determined by the more contractile cells within the doublets. These results thus revealed that contractile leader cells could drive the chiral motion of minimal collectives.

**Significance Statement:** Chirality, which represents a fundamental property of living systems, manifests in cell collectives by their persistent biased directional swirling. Despite the clear identification of the implication of actomyosin cytoskeleton in driving the internal chiral symmetry break occurring in cells, little is known about the actual role of cellular forces produced by this network in the development of handedness in collectives. Our findings establish that the level of mechanical energy developed by pairs of confined endothelial cells regulates the strength and direction of their rotation. Our results also identify the more contractile cell of the doublet as the cell driving the direction and speed of rotation of the pair. This study thus sheds new light on the importance of the generation and integration of mechanical forces within a small collective in the determination of its chiral rotation.

## Introduction

Chirality, defined as the property of an object not to be superimposable to its mirror image, is an evolutionary-conserved characteristic of living organisms (1, 2). Chirality notably manifests in left– right (LR) asymmetry of tissues during embryonic development (3–5), where increasing evidence suggests that the left-right bias of their constituting cells significantly contributes to this process (6, 7).

At the cellular level, chirality has been shown to manifest in various forms (8), among which collective cellular biased motion represents one of the most persistent depictions, likely due to the development of long-range correlations between individual cells (9). In most of the confined eukaryotic cells grown as collectives, a biased cellular alignment with respect to the system boundaries is indeed observed (10–13) and is eventually associated with the persistent chiral swirling of cells along the boundaries in both 2D (14, 15) and 3D systems (16, 17).

Although the LR bias of cells has been shown to strongly rely on the activity of several groups of actin-related proteins (5, 7, 10, 18, 19), the conditions leading to the emergence of chirality in motile multicellular groups is far from being understood. According to several studies, the presence of an intact actomyosin network notably appears as a key determinant of cellular chirality expression. Myosin activity has indeed been shown to be indispensable for both the chiral symmetry breaking of the actin cytoskeleton in single cells (20–22) and the biased alignment of cell ensembles (11), for which a LR-polarized distribution of myosin II is further described as concomitant with the appearance of cell chirality in the myocardium tissue (7). Thus, although cell mechanics appear to be a key factor that can affect the chiral swirling observed in cell collectives, the exact role of cellular forces in chiral bias expression remains to be determined.

Importantly, what leads to the expression of opposite handedness among different cell types sharing a priori common chiral actomyosin template remains unclear (14). These variations first reflect the dependence of the chirality manifestation on additional unknown factors that are sensitive to environmental conditions (23) such as the size (24, 25) and geometry of the confinement (26), or the rigidity of the substrate on which cells are evolving (27). Second, although the chiral manifestation has been well established in big collectives, the constituting cells display a complex and heterogeneous behavior due to the local environment they interact with. This is well illustrated by the different morphologies or alignment cells adopt along the width of rectangles or rings (10, 28) and the counter-rotations they display on geometries exhibiting several boundaries with opposite curvatures (14). As a result, this inter-cellular variability interferes with the behavior of the collective and renders difficult the identification of the critical parameters involved in the chiral phenotype expression within multicellular systems.

In this study, we first aimed at reducing the number of cells in a moving collective in order to identify the minimal set of conditions allowing the emergence of a chiral movement. We then combined live monitoring of cell movement with traction force microscopy to investigate the role of contractile forces in the establishment of the chiral bias.

## Results

### HUVEC doublets represent a minimal cell collective system displaying chirality

Several studies have established that endothelial cells of various origins display a robust chiral bias, manifested by their biased alignment and/or chiral swirling when grown as collectives in confined environments (12, 14, 21, 29, 30). More recently, the chirality of endothelial cells was also demonstrated *in vivo* in vessels (13). Importantly, in both systems, chirality plays a crucial role in the modulation of the endothelium and vascular permeability (12, 13), highlighting its functional role in the physiology of these tissues. In this context, we selected endothelial cells as a model collective system in our study. First we sought to characterize their chiral bias in a simplified and controlled environment. To this end, we confined the cells on disk-shaped micropatterns, which present the advantage of displaying a plain 2D surface with only one external boundary, thereby decreasing the complexity of the geometries tested in previous studies (ring micropatterns or tubes). We plated HUVECs at high-density on 250 µm diameter adhesive disks and imaged them for 8 to 14 hours (Fig. 1A and S1A). By characterizing the rotation in these systems within this time window, we identified two classes of collectives: collectives in which the motion of individual cells displayed no coherence (designated by “no coherent rotation” or “NC”) and those where the majority of cells aligned and moved in a coordinated fashion, essentially along the periphery of the disk (Fig. 1A). On average, 60% of rotating cells demonstrated a coherent rotation, while 40% displayed no preferential orientation (Fig. 1B), showing that the majority of cells constituting the collectives were able to rapidly reorganize and polarize along the external boundary, ultimately leading to their coordinated motion (Fig. S1B and Movie 1). To test the impact of the initial cell density on the coherence of the motion, we counted the initial number of nuclei per collectives for the two subpopulations and found no significant difference between them (Fig. S1C), indicating that the initial confluence of cell ensembles played a minimal role in triggering their coherent motion. Interestingly, around 90% of the collectives rotating in a coordinated fashion displayed a CW bias (Fig. 1C and Movie 2), in agreement with previous works (14). We next analyzed the speed of rotation of the nuclei located in different zones of the collective: along the external boundary in a radius of 70 µm (termed “OUT”) and in their inner part (termed “IN”) (Fig. S1D and E). We observed that cells at the periphery of the collectives moved more coherently and with a significantly higher speed than those in the internal part. These results indicate that even in the absence of an internal boundary, big collectives still display a high heterogeneity in their rotational behavior. Consequently, we wondered whether we could simplify the endothelial collective to a more minimalistic configuration while maintaining the expression of the chiral phenotype. To do so, we gradually reduced the number of HUVEC cells and examined the rotation of clusters constituted of either 4 (quadruplets), 3 (triplets) or 2 cells (doublets) confined on disks of 60 µm diameter (Fig. S1A and 1D). In all these cases, the percentage of minimal collectives rotating persistently reached about 80% to 90% (Fig. 1E), which was significantly higher than the 60 % rotation measured in the case of bigger collectives over the same period. This result suggests that reducing the number of cells constituting the collective tends to normalize the environment encountered by individual cells thereby favoring their coherent motion. These data are in good agreement with previous work showing that the persistence of coherent angular motion within a cluster of cells exhibits discontinuity above four cells and further requires their geometric rearrangement (31). Importantly, our quantification of the rotation direction among the population of persistently swirling cells showed that the CW-bias measured in big ensembles was maintained in the minimal collectives tested (Fig. 1F and Movie 3). In quadruplets, the CW-bias was lower than that in large collectives and was clearly visible only in 75% of the rotating minimal collectives. In triplets and doublets, the CW-bias was further decreased, reaching about 60% of the rotating collectives. However, this proportion turned out to be extremely robust within independent experiments (N=16 experiments; n=2909 cells) and stable overtime (up to 39 hours) (Fig.1 SF-J and Movie 4). Overall, these results demonstrate that the chiral bias displayed by big cell collectives is reduced but conserved among cell doublets, in which the coordinated rotation was also more frequent. We thus decided to focus our study on endothelial cell doublets since their simplicity, in addition to their robust chiral bias offer a unique opportunity to investigate the critical parameters determining the CW or CCW directionality of the collective rotational motion.

**Figure 1.**
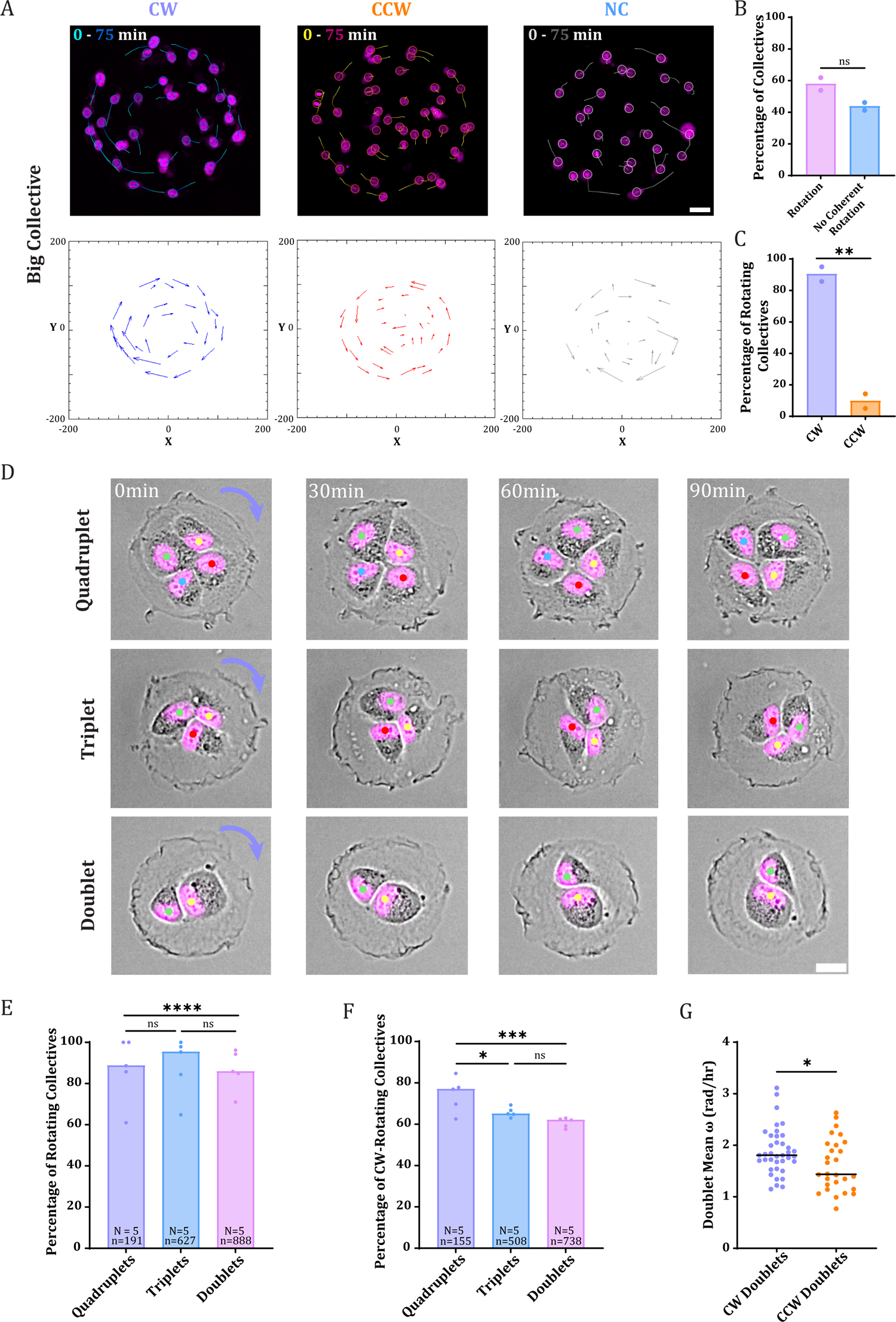
Minimal endothelial collectives exhibit a weak but robust clockwise bias. A. Upper panel: single time-point images of HUVEC cell collectives on disk-shaped micropatterns (250 µm diameter) rotating either coherently to the right (CW) or to the left (CCW) or not rotating coherently (NC). Nuclei are stained with Hoechst (magenta) and trajectories determined in TrackMate over 5 frames are represented by tracks on the images (the circles around the nuclei indicate the last time point of the trajectories). Lower panel: vector plot of the trajectories generated in Fiji. Scale bar=30µm. B. Repartition of the percentage of coherently rotating collectives (Rotation) and non-coordinated ones (No coherent Rotation). N=2 independent experiments. n=172 collectives. Statistical significance was assessed using unpaired t-test (p=0.0948.). C. Repartition of the percentage of rotating minimal collectives between CW and CCW-rotation. Statistical significance was assessed using unpaired t-test (p=0.0066). D. Time-lapse series of cells and nuclei (Hoechst in magenta) within collectives of decreasing complexity (top panel: quadruplet; middle panel: triplet; bottom panel: doublet). Each nucleus is marked by a different colored spot and is identified overtime in the sequence. The purple arrow indicates the CW direction of rotation of the collective. Scale bar=15µm. E. Percentage of rotating collectives. N=5 independent experiments. Statistical significance was assessed using Chi-square test (Fischer’s exact; ns_Quadruplets-Triplets_>0.9999; ns_Triplets-Doublets_=0.3063; ****<0.0001). F. Percentage of CW-rotating minimal collectives. Statistical significance was assessed using Chi-square test (Fischer’s exact; Significance testing: ns=0.6604; *=0.0133; ***=0.0002). G. Mean angular velocities of the doublets rotating CW and CCW. N=1; n=66 doublets. Statistical significance was assessed using an unpaired t-test (p=0.0256).

### The stored energy of the doublets correlates to their swirling speed and direction

Interestingly, when measuring the angular velocities of the doublets to characterize their swirling (Fig. 1G), we found that their speed of rotation was significantly lower when going in the CCW direction than in the CW one. In the context of cell migration, several studies previously demonstrated that both the speed and persistence of migration are dependent on cellular contractility levels (32–35). We therefore tested next whether CW- and CCW-rotating doublets could exhibit different contractile properties. To this end, we plated HUVEC cells on 60µm disk-shaped micropatterns fabricated on polyacrylamide gels of 17 kPa and performed traction force measurements over the course of rotation (Fig. 2A and Fig. S2A). This method allowed us to compute the stored mechanical energy of the doublets overtime (Fig. 2B, Fig. S2B and Movie 5). By additionally contouring the junction, we could also measure the doublet angular velocity over time (Fig. 2C). By averaging the stored energy of distinct rotating doublets (Fig. 2D), we first noticed that the mean ME of the CCW-rotating doublets was significantly higher than that of the CW-rotating pairs of cells (Fig. 2D and 2E). This was also accompanied by a significant decrease in the average length of the junction in CCW-rotating cells (Fig. S2C), in line with previous works showing that junction length depends on actomyosin contractility (36, 37). We then assessed whether this difference in the contractility level of the doublets could also be associated with a variation in their rotation speed. When plotting the angular velocity of the CW- and CCW-rotating doublets averaged overtime (Fig. 2F), we first observed that the angular velocity was lower in CCW-rotating cells compared to CW-rotating doublets, in line with our previous measurements on glass (Fig. 1G). Finally, plotting the mean angular velocity of doublets against the corresponding magnitudes of mean stored ME revealed a noisy but significant negative correlation between these two parameters (Fig. 2G). Together, these results demonstrate the existence of a correlation between the contractility level of the doublets and the speed and direction of their rotation.

**Figure 2.**
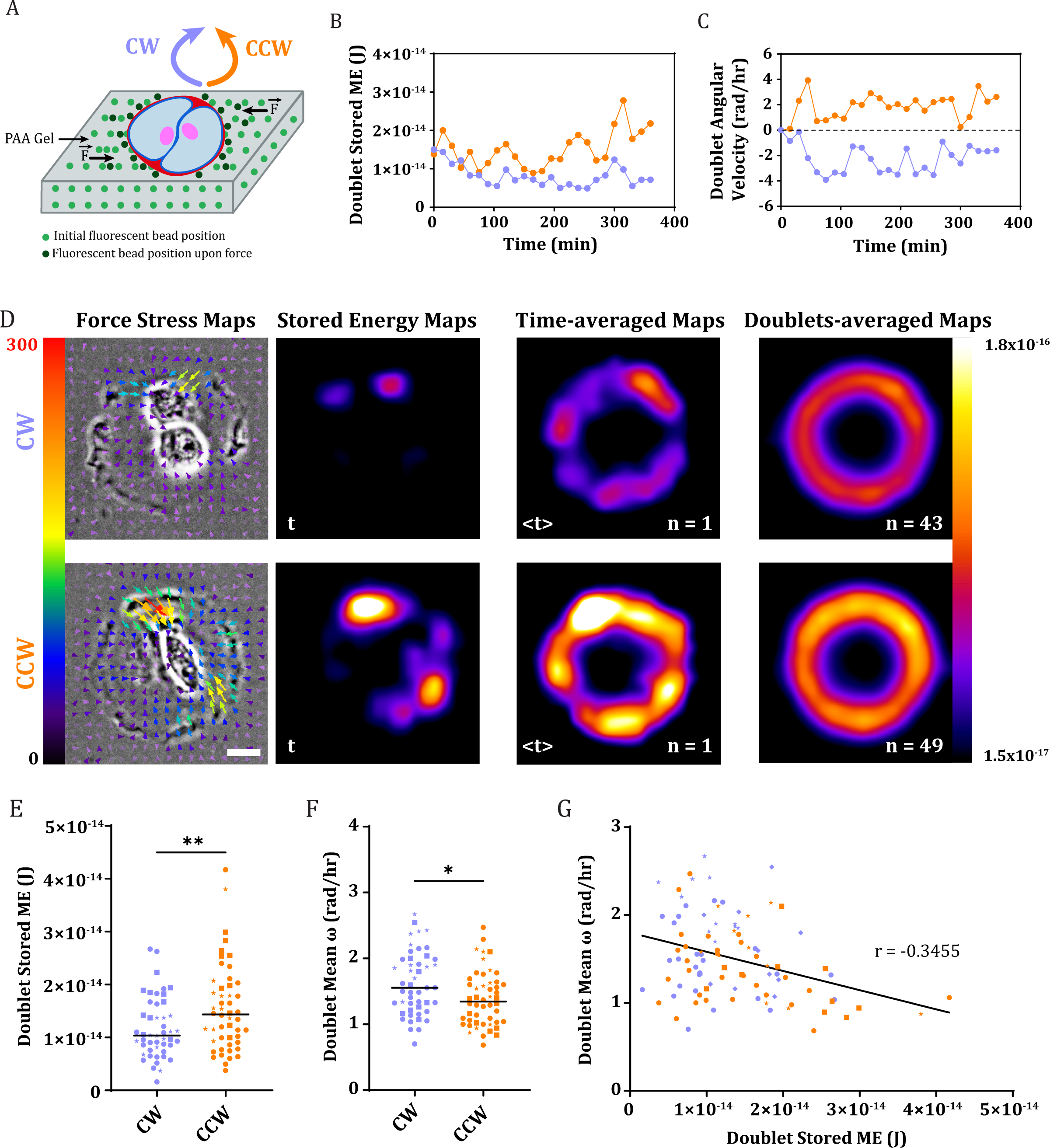
The stored ME of pairs of endothelial cells correlates to their chiral rotation. A. Schematic representation of the experimental setup used for TFM. Cell doublets (blue) were seeded on disk-shaped micropatterns covered with a mix of collagen and fibronectin (red). While rotating, the doublet exerts forces F on the substrate, leading to the displacement of the beads (green dots) embedded in the gel. B. Temporal profile of the stored mechanical energy (ME) of a doublet rotating CW (purple) and a doublet rotating CCW (orange). C. Temporal profile of the instantaneous angular velocity of a doublet rotating CW (purple) and a doublet rotating CCW (orange). D. From left to right: Force stress map overlaid onto phase contrast images of a CW (top) and a CCW (bottom) rotating doublet at a single time point. Traction force scale bar in Pa. Image scale bar=15µm. Corresponding Stored Energy maps (second column). Representation of the stored energy for the corresponding doublets averaged overtime (third column; n=1). Averaged Stored Energy Maps of distinct CW (above) and CCW (below) rotating doublets (n=43 and 49 respectively). Color-coded scale bar in J. E. Mean stored ME of doublets computed over their CW and CCW rotation. Statistical significance was assessed using an unpaired t-test (p=0.0087). N=3 independent experiments. n=98 doublets. F. Mean angular velocity of CW and CCW rotating doublets. Statistical significance was assessed using an unpaired t-test (p=0.0110). N=3 independent experiments. n=98 doublets. G. Mean angular velocity of the rotating doublets as a function of their corresponding mean stored ME. CW rotating doublets are represented in purple and CCW rotating doublets in orange. N=3 independent experiments. n=98 doublets. The Pearson correlation coefficient r is indicated on the plot. A linear regression fit was applied on the data.

### The level of contractility of the doublets modulates the directionality of the chiral bias

To test whether variations of the forces developed by the doublets could affect the expression of the chiral bias of the minimal collective, we further used chemical inhibitors to instantaneously modulate the contractility of the doublets.

We first treated the doublets with different concentrations of Rho kinase inhibitor (ROCKI) shortly after seeding the cells on micropatterns to decrease their contractility. As expected, we observed a reduction in the formation of actomyosin contractile fibers, characterized by a significant decrease in phosphorylated myosin light chain (p-MLC) upon treatment compared to the control conditions (Fig. 3A). These changes were also associated with a noticeable drop in the traction forces measured within the same doublets (Fig. 3B). We then monitored the doublets over time in the presence of increasing concentrations of ROCKI and quantified the percentage of rotation among the doublets (Fig. 3C and Movie 6). A gradual and significant decrease in the percentage of rotating doublets was registered upon increasing the inhibitor concentration, indicating that a minimal contractility level was required for the onset of rotation. This result was accompanied by a significant drop in the mean angular velocity of the doublets in the presence of the inhibitor at 7µM (Fig. 3D). Importantly, we recorded a slight but significant increase in the percentage of CW-rotating doublets with increasing concentrations of ROCKI (Fig. 3E). The same trend was also observed in response to increasing concentrations of Blebbistatin (Fig. S3A-C). These results supported the hypothesis that low mechanical energies favor a CW-chiral bias in doublet rotation.

**Figure 3.**
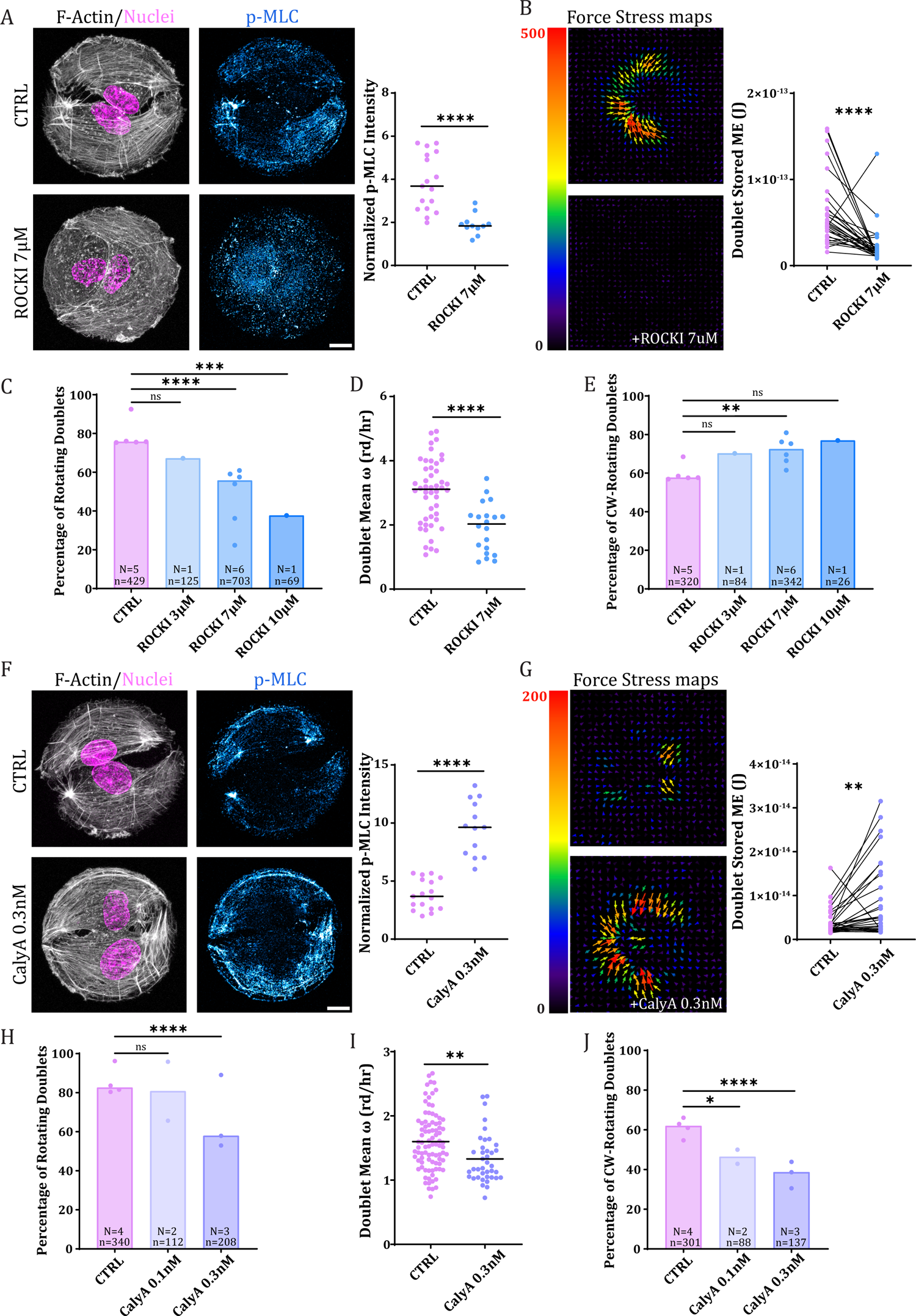
The level of contractility of the doublets modulates the directionality of their chiral bias. A. Left panel: Immunostaining of control (top panels) and Rho kinase inhibitor (ROCKI)-treated doublets (7µM for 6hrs; bottom panels). Stainings: F-actin (phalloidin), nuclei (Hoechst), and phospho-myosin (p-MLC). Right panel: quantification of the p-MLC intensity signal for control and ROCKI-treated doublets after normalization by the fluorescence of fiduciary markers. N=1; n= 28 doublets. Statistical significance was assessed using an unpaired t-test (p<0.0001). Scale bar=15µm. B. Left panels: traction stress map of a doublet before (top image) and after treatment with ROCKI at 7µM (bottom image). Force scale color bar in Pa. Right panel: stored ME measured in the same doublets before and after treatment with ROCKI. N=1; n=30 doublets. Statistical significance was assessed using a paired t-test (p<0.0001). C. Percentage of rotation quantified in control doublets and in doublets treated with increasing concentrations of ROCKI (3, 7 and 10µM). N indicates the number of individual experiments and n the total number of doublets used for quantifications. Statistical significance was assessed using Chi-square test (Fischer’s exact; Significance testing: ns=0.1099; ***=0.0007; ****<0.0001). D. Mean angular velocity of control and ROCKI-treated doublets. Statistical significance was assessed using an unpaired t-test (p<0.0001). N=1. n=68 cells. E. Percentage of CW-rotating doublets in control condition and in the presence of increasing concentrations of ROCKI (3, 7 and 10µM). N indicates the number of individual experiments and n the total number of doublets used for quantifications. Statistical significance was assessed using Chi-square test (Fischer’s exact; Significance testing: ns_CTRL-RockI3µM_=0. 1282; ns_CTRL-RockI10µM_=0.1394; **=0.0087). F. Left panel: Immunostaining of control (top panels) and CalyA-treated doublets (0.3nM for 6hrs; bottom panels). Stainings: F-actin (phalloidin), nuclei (Hoechst) and phosphor-myosin (p-MLC). Right panel: quantification of the p-MLC intensity signal for controls and CalyA-treated doublets after normalization by the fluorescence of fiduciary markers. N=1; n=30 cells. Statistical significance was assessed using an unpaired t-test (p<0.0001). Scale bar=15µm. G. Left panels: traction stress map of a doublet before (top image) and after treatment with CalyA at 0.3nM (bottom image). Force scale color bar in Pa. Right panel: stored ME measured in the same doublets before and after treatment with CalyA. N=1; n=30 doublets. Statistical significance was assessed using a paired t-test (p=0.0029). H. Percentage of rotation quantified in control doublets and in doublets treated with increasing concentrations of CalyA (0.1 and 0.3nM). N indicates the number of individual experiments and n the total number of doublets used for quantifications. Statistical significance was assessed using Chi-square test (Fischer’s exact; Significance testing: ns=0.0115; ****<0.0001). I. Mean angular velocity of control and CalyA-treated doublets. Statistical significance was assessed using an unpaired t-test (p=0.0014). N=1. n=128 doublets. J. Percentage of CW-rotating doublets in control condition and in the presence of increasing concentrations of CalyA (0.1 and 0.3nM). N indicates the number of individual experiments and n the total number of doublets used for quantifications. Statistical significance was assessed using Chi-square test (Fischer’s exact; Significance testing: *=0.01238; ****<0.0001).

Using Calyculin A (CalyA), an inhibitor of protein phosphatase 2A and 1, we then tested the effects of a force increase on the rotational chiral bias. As expected, low doses of CalyA induced a reinforcement of contractile actomyosin fibers in the doublets and significantly increased the amount of p-MLC compared to control conditions (Fig. 3F). This phenotype was also associated with a significant increase in traction stresses exerted by the doublets (Fig. 3G). Interestingly, low doses of CalyA led to a significant decrease in the percentage of rotating doublets (Fig. 3H, Movie 7) and reduced their angular velocity (Fig. 3I), showing that an excess in contractility impairs the rotation. More interestingly, the percentage of CW-rotating doublets gradually decreased with increasing CalyA concentrations (Fig. 3J).

Taken together, these results first show that an intermediate level of contractility is required to induce doublet rotation. This biphasic behavior is consistent with previous studies of showing the existence of an optimal adhesion strength and contractility in the regulation of single cell migration speed (32, 38, 39). More importantly, our results also demonstrate that low doublet contraction favors a CW bias whereas high contractility induces a CCW bias.

### The mechanical energy of the more contractile cell of the doublet is a good predictor of its chiral rotation

When considering collective motion, the magnitude and distribution of the forces exerted by individual cells can affect the behavior of the collective (40, 41). This led us to examine the mechanical role of each cell within the doublets in the determination of their chiral bias.

In order to isolate the contractile forces of individual cells within the rotating doublets, we contoured each cell of the pairs and computed their individual stored energy over time (Fig. 4A). Despite noticeable fluctuations, we noted that on average one of the two cells was slightly more contractile than the other. To visualize this asymmetry, we rotated the traction maps of each doublet such that the cell displaying the higher ME, hereon referred to as the Stronger cell, would lie to the right side of the map and averaged them overtime (Movie 8). We then overlaid these averaged rotated maps for distinct doublets (Fig. 4B). This representation clearly highlighted the existence of an asymmetry in the mean stored ME between the two cells constituting the doublets, which was visible in both types of rotating pairs. To characterize this asymmetry between doublets of opposing directionalities, we then plotted the ratio of stored ME (Stronger cell/Weaker cell) between the two cells of the doublets for CW- and CCW-rotating pairs (Fig. 4C). These data showed no significant difference between these two sub-populations, demonstrating that the magnitude of the asymmetry in contractile energy between the two cells of the doublet was not correlated to the doublet bias. To challenge the mechanical impact of individual cells constituting the doublet, we next compared the stored ME of cells exhibiting low contractile forces (Weaker cells) and high contractile forces (Stronger cells) in CW and CCW-rotating doublets (Fig. 4D). We first noticed that Weaker cells within CW or CCW-rotating doublets displayed similar stored ME levels. By contrast, Stronger cells exhibited significantly higher ME when they were part of CCW rotating doublets. Additionally, the mean angular velocity of doublets rotation was negatively correlated to the ME of Stronger cells (Fig. 4E; Pearson=-0.36), as observed in the case of doublets stored ME (Fig. 2G). These results suggested that the ME of Stronger cells could be associated with the chiral bias of the doublets. To test this hypothesis, we then calculated the relative frequency of Stronger cells to be part of a CW or CCW-rotating doublet, depending on the level of their contractile energy (Fig. 4F). Importantly, we noticed that the relative frequency of CW directionality was high for lower values of stored ME and decreased gradually upon increased contractility of Stronger cells. On the contrary, the frequency of CCW rotation, initially low at reduced contractility levels, progressively increased with increasing magnitudes of stored ME of the Stronger cells. By contrast, both the angular velocities and the relative frequency of the direction of rotation were poorly correlated to the mechanical state of the Weaker cell (Fig. S4A and 4B). Altogether, these findings put forward that the mechanical energy of Stronger cells within the doublets can reliably predict their rotational bias.

**Figure 4.**
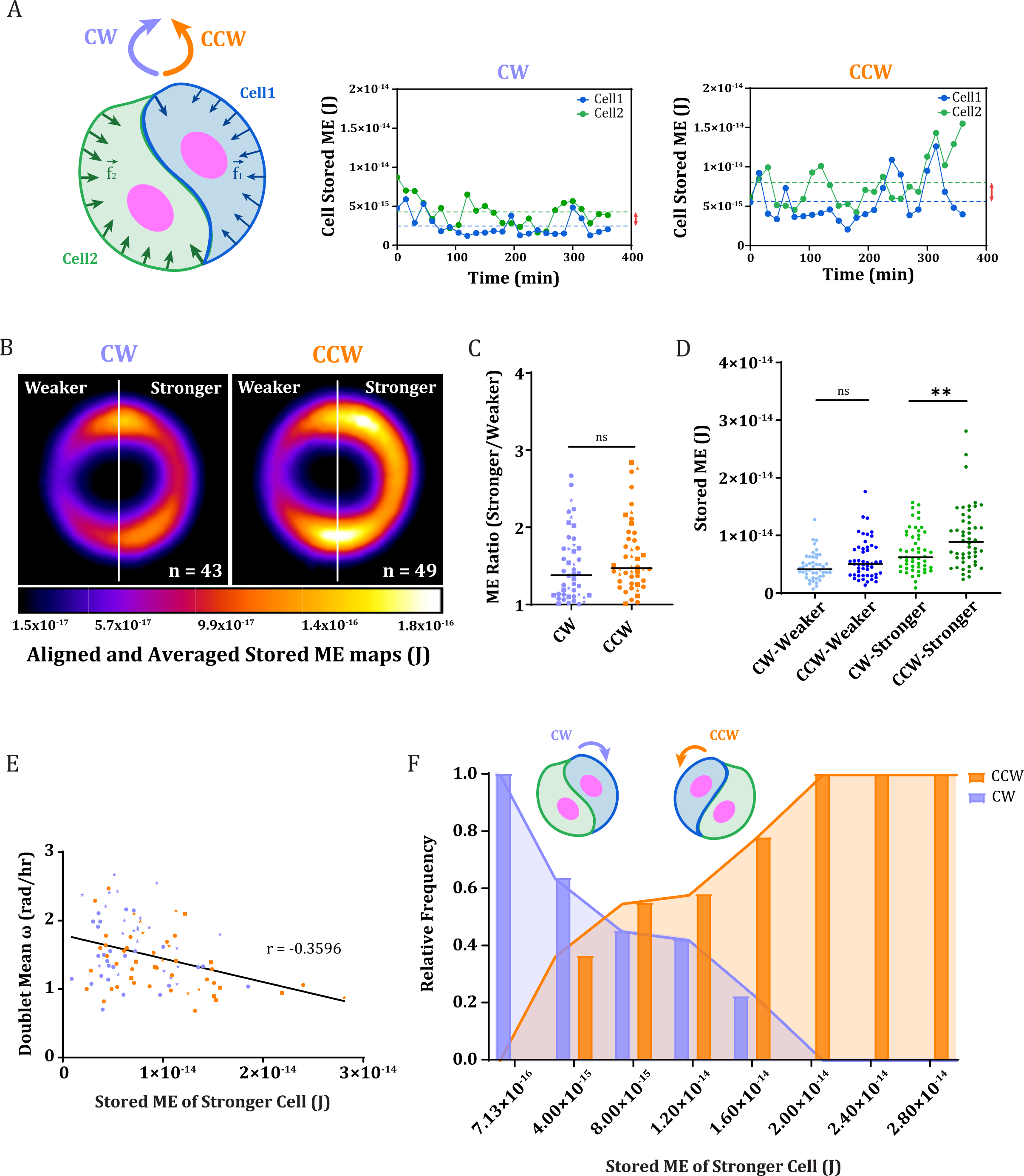
The mechanical energy of the more contractile cell of the doublet is a good predictor of its chiral rotation. A. Left panel: Schematic representation of the repartition of the tractions forces exerted by individual cells constituting a doublet. Right panel: temporal profiles of the stored ME of individual cells constituting a CW-rotating doublet (left plot) and a CCW-rotating doublet (right plot). The mean difference in stored ME is highlighted by the red double arrow. B. Averaged representation of the stored ME maps of distinct CW (left image) and CCW-rotating doublets (right image) after their alignment with the stronger cell being on the right side of the image. The number of maps averaged is indicated on the images. Color coded scale bar in J. C. Comparison of the ME ratio of the Stronger cell/Weaker cell between CW- and CCW-rotating doublets. Statistical significance was assessed using an unpaired t-test (p=0.2038). N=3 independent experiments. n=98 doublets. D. Comparison of the Mean stored ME of individual cells (weaker and stronger) constituting CW- or CCW-rotating doublets. Statistical significance was assessed using an unpaired t-test (Significance testing: p_weaker_=0.1145; p_stronger_=0.0050). N=3 independent experiments. n_CW_=94 cells and n_CCW_=100 cells; n=98 doublets. E. Mean angular velocity of the doublets as a function of the Stored ME of the corresponding stronger cells. The Pearson correlation coefficient is represented on the graph. N=3 independent experiments; n=98 doublets. A linear regression fit was applied on the data. F. Relative frequency distribution of CW (purple) and CCW (orange) rotating doublets as a function the stored ME of their stronger constituting cell. Frequencies were calculated on n_CW_=47 doublets; n_CCW_=50 doublets.

### The more contractile cell of the doublet drives the chiral bias of the pair

We then wondered whether the more contractile cell of the doublet could actually drive the rotational bias of the doublet. To that end, we decided to modulate the contractility of only one cell of the doublet by using a different cell type than HUVECs.

We first tested if MEFs, predicted to be more contractile than HUVECs (42) and displaying a more prominent actomyosin network than HUVECs (Fig. S5A-C), were indeed exerting higher traction forces than endothelial cells. To that end, we performed TFM measurements in MEFs (Fig. S5D) and HUVECs homotypic doublets (Fig. 5A) using a common experimental setup (60 µm disks and 17kPa PAA gels). As expected, the results showed that MEF doublets developed significantly higher ME than HUVEC ones (Fig. 5B). To test the impact of such a higher contractility level of MEF homotypic doublets compared to HUVECs, we next used live imaging to assess the rotation speed and bias of both types of doublets. We found that MEF doublets rotation was less frequent (Fig. 5D) and slower than HUVEC doublets (Fig. S5E). Interestingly, homotypic MEF doublets exhibited a predominance for CCW rotation (60%, n=434) unlike HUVEC cells that displayed the expected CW-bias (60%, n=358) (Fig. 5E). This further confirmed our conclusion that high cell contractility favors a CCW rotation.

**Figure 5.**
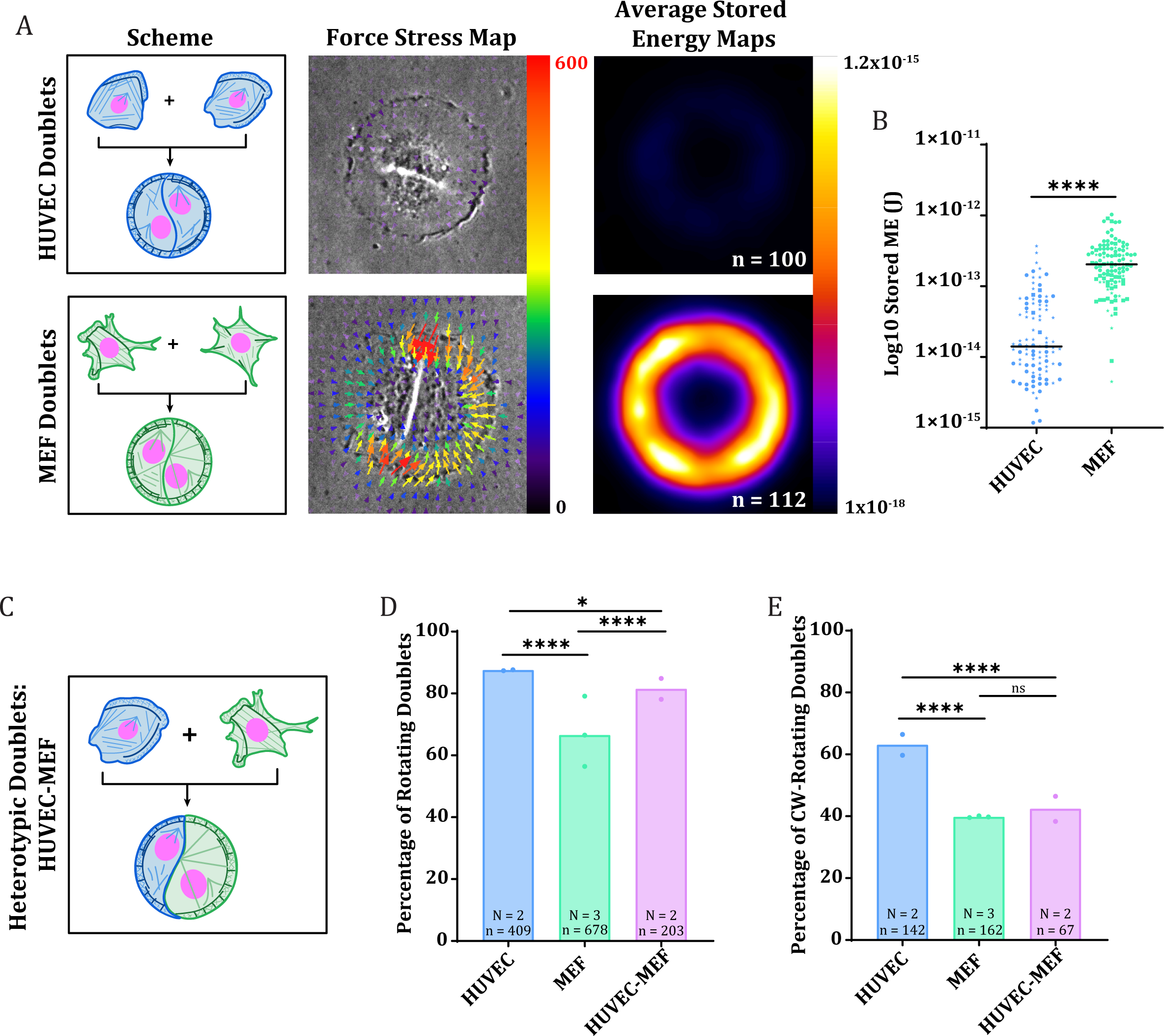
The stronger contractile cell among the pair drives the chiral rotation of the doublet. A. From left to right: Schematic representation of HUVECs and MEFs constituting homotypic doublets. Force stress maps overlaid onto phase contrast images of a HUVEC homotypic doublet (top image) and a MEF homotypic doublet (bottom image) at a single time point. Traction force scale bar in Pa. Averaged Stored Energy maps of distinct doublets. The number of maps averaged per condition are indicated on the images. Color coded scale bar in J. B. Stored ME of homotypic HUVEC and MEF doublets. N=3 independent experiments; n=100 doublets for HUVEC and 112 for MEF. Statistical significance was assessed using unpaired t-test (p<0.0001). C. Schematic representation of HUVECs and MEFs constituting heterotypic doublets. D. Percentage of rotating doublets in the three populations. The number of independent experiments (N) and total number of cells analyzed (n) are indicated on the graph. Statistical significance was assessed using Chi-square test (Fischer’s exact; Significance testing: *=0.0217; ****<0.0001). E. Percentage of CW-rotating doublets in the three populations. The number of independent experiments (N) and total number of doublets analyzed (n) are indicated on the graph. Statistical significance was assessed using Chi-square test (Fischer’s exact; Significance testing: ns=0.7796; ****<0.0001).

To assess the effect of the more contractile cell on the chiral rotation, we then examined the rotation of heterotypic systems, in which doublets were composed of a less contractile cell, a HUVEC with a CW-bias, and a more contractile cell, a MEF displaying a CCW-bias (Fig. 5C, and 5D). We pre-labelled MEFs with Calcein-AM to clearly identify the heterotypic doublets. By monitoring heterotypic doublets over several hours, we noticed that, on the contrary to HUVECs and MEFs homotypic doublets for which the individual cells often occupied a comparable space on the micropattern, MEFs occupied most of the space on the disk (Fig. S5F). HUVECs were confined to the periphery of the doublets along which they moved more than the central MEF (Movie 9). In addition, heterotypic doublets exhibited an intermediate rotation percentage (70%, Fig. 5D) compared to both HUVEC (90% of rotation) and MEF (60% of rotation) homotypic doublets. Interestingly, the chiral bias of heterotypic doublets was CCW (60%, n=163) as in the case of MEF doublets (Fig. 5E). This result demonstrated that the more contractile cell of the doublets could drive the motion of the heterotypic system.

## Discussion

Our findings first reveal the link between the chiral bias of migrating cells and their contractile energy, supporting the idea that cell chirality is a mechanosensitive process relying on actomyosin contractility (11, 27). Indeed previous work reported the loss of chiral bias upon a disruption of the actomyosin network (20, 27, 43). In addition, the localization of myosins was found to be correlated with the orientation of the chiral bias in cell doublets (17) and in tissues (7, 44). In this study, by directly measuring both the forces and the speed of migrating cells, and by modulating these forces over a broad range with chemical drugs, we demonstrated that the chiral bias is determined by the magnitude of contractile forces. In the endothelial and fibroblasts cells we tested, high forces appear to promote CCW bias, whereas low forces tend to impose a CW bias. Whether this relationship can be generalized to other cell types would require further investigation. However, it is interesting to hypothesize that the cellular contractility levels determines the corresponding chiral bias of their tissues.

In addition, the analysis of the mechanical energy of individual cells within the doublet consistently revealed significant differences between the two cells. In bigger cell ensembles, a similar heterogeneity and force imbalance was previously described to impact polarization and migration of the collective with follower cells pulling on the future leader cell (40). Interestingly, the emergence of a leader cell, that ensures front–rear polarization and guidance among other cells via mechanical signaling, is associated with the exertion of higher traction forces (45). We found that the more contractile cell with of the doublet imposes its own bias on the other. Since a mechanical coordination can happen by the transmission of contractile forces between adjacent cells (46), it is tempting to speculate that the biased distribution of these forces can also be transmitted and impact the chirality of the collective. These data overall provide additional support to the hypothesis that chiral organization in multicellular groups is determined by the chirality developed in individual cells (6, 10, 47, 48). However, despite numerous studies on the migration of individual cells, only a few reported the spontaneous development of chiral bias in cell trajectories (49–51). This suggests that it is the integration of individual cells in a moving collective that allows their dominant bias to be revealed.

Because the chiral manifestation in individual cells is rare and transient (20), its underlying mechanism remains unclear. Several studies suggest that a combination of cytoskeleton and motor proteins can generate an individual cellular torque (4, 19, 20, 25). The imbalance between radial fibers polymerization, executed by formins, and myosin-driven contractility of transverse arcs has been proposed to control the direction of the chiral bias (20, 52). According to this hypothesis, myosin II activity would regulate the strength of the CW bias by modulating the prominence and integrity of transverse arcs at the expanse of radial fibers. Our results are consistent with this hypothesis, but the exact role of the migration process and of the integration in the collective requires further investigation. Taken together these studies point at the need to study not only the architectures, but also the intracellular displacement and dynamics of networks to understand the propagation of chiral bias from the right-handed actin filaments to the individual cell and the chiral bias of multicellular collectives.

## Materials and Methods

### Cell Culture

Human Umbilical Vein Endothelial Cells (HUVEC-h-TERT2 – Evercyte CHT-006-0008) were cultured in Endothelial Cell Growth Medium (Lonza EGM BulletKit CC-3121 & CC-4133) supplemented with the growth factors provided in the kit, 2% fetal bovine serum, and 1% antibiotic-antimycotic solution (Gibco 15240062). The culture flasks were coated with Gelatin (Sigma G1890) 0.1% in Dulbecco’s phosphate-buffered saline (DPBS Gibco 14200075) for 30 minutes before use. Mouse Embryonic Fibroblasts (MEFs) originating from Goldman’s lab (53) were cultured at 37°C and 5% CO2 in DMEM supplemented with 15% fetal bovine serum and 1% antibiotic-antimycotic solution. The cells were grown at 37°C and 5% CO2 and tested regularly for Mycoplasma using VenorGeM Advance (11–7024) PCR kit. When passaging, the cells were washed once with DPBS (Gibco 14200075) and detached using TrypLE Express Enzyme (Gibco 12605010).

### Micropatterned Glass Coverslips (CS) Preparation

Protein micropatterns on glass coverslips were prepared as previously described in the literature (54) following the steps described below.

Cleaning

CS (EPREDIA) 20×20mm or 25×25mm #1.5 (CS) were submitted to three rounds of washing: sonication in acetone (Carlo Erba 528203) for 30mins, sonication in isopropanol (Sigma 34863) for 30mins, and sonication in MilliQ water for 30mins. CS were then air-dried.

#### Polystyrene Coating

To promote cellular adhesion, the CS were coated with a thin layer of polystyrene on top of an adhesion promoter. Clean CS were activated by an air plasma treatment (Diener electronic GmbH & Co KG - Zepto) then placed inside a semi-closed container heated at 75°C in the presence of a few drops of Hexamethyldisilazane (HMDS Sigma 440191) for a least 6hrs. Alternatively, a layer of Ti-Prime was spin-coated (Laurel – WS-400BBX-6NPP) on the CS for 30secs at 1000 rpm. The CS were then placed in the spin coater (Laurell Technologies), covered with a solution of Polystyrene (PS) 1% (Acros Organics AC404720250) in toluene (108325), and spun for 30secs at 1500 rpm.

#### PLL-PEG Coating – Passivation

The PS-coated CS were activated by air plasma treatment and then coated with PLL-PEG solution (PLL-PEG JenKemTechnology ZL187P072) 0.1mg/mL in HEPES (H3375) 10mM, pH 7.4 in MilliQ water for 30mins at room temperature. After incubation, the CS were dewetted using HEPES 10mM and stored at 4°C for at least 1 hour before use.

#### Deep UV Micropatterning

PLL-PEG-coated CS were placed on a vacuum holder in tight contact with a quartz-chrome printed photomask (Toppan Photomask). The sandwich was then placed in a pre-warmed UVO cleaner (Jelight - Model No. 342A-220), with a power of 6mW/cm2, for 5 min, during which the PLL-PEG layer was burned with deep UV (190nm). After exposure, the patterned CS were gently detached from the mask using vacuum and coated with the protein solution.

#### Protein Coating

The patterned CS (from the previous step) were coated with a protein solution, composed of fibronectin (FN Sigma F1141) 20µg/mL ± Fibrinogen from Human Plasma, Alexa Fluor 546 or 647 Conjugate (FNG Invitrogen F13192 or F35200) 10µg/mL in sodium bicarbonate (NaHCO3 Sigma S6297) 100mM in MilliQ water for 30mins protected from light. After incubation, the patterned CS were washed 3 times with NaHCO3 and dewetted. Before seeding the cells, the CS were washed with sterile DPBS.

### Polyacrylamide Gel Micropatterning for TFM

Patterned hydrogels were prepared according to the “Glass Method” previously described in (55). Briefly, 22×22mm #1.5 glass CS were passivated with PLL-PEG, deep UV-patterned, and coated with adhesive proteins (FN 20µg/mL or a mixture of FN and Col (Collagen I, rat tail Gibco A1048301) 10:10µg/mL) as previously described. A mixture of 40% Acrylamide (Sigma A4058) and 2% Bis-acrylamide (Sigma M1533) in MilliQ water (experimental Young modulus of 16.7kPa) was prepared according to (56), degassed in a vacuum bell during protein incubation, mixed with the passivated 200nm fluorescent beads (FluoSpheres Carboxylate-Modified 200nm Polystyrene Microspheres Invitrogen F8810 – 580/605nm or F8807 – 660/680 nm), sonicated for homogenization, and kept on ice. 1 µL of Ammonium persulfate (APS Sigma A3678) and 1 µL of N,N,N′,N′-Tetramethylethylenediamine (TEMED T9281) were added to the 165 µL PAA mix. 25µL of the latter solution was added on each of the 22×22 mm patterned CS. Silanized 20×20 mm CS were flipped on top of the PAA drops, and the gel was allowed to polymerize for 40 to 50mins. At the end of the incubation, the sandwiched gels were submerged with NaHCO3 for a few minutes before they were gently detached from the patterned CS using a scalpel. Micropatterned PAA gels were then stored in 35mm Petri plates submerged in NaHCO3 at 4°C until use. Before seeding the cells, the gels were washed twice with sterile DPBS and once with warm medium.

### Cells seeding on micropatterns

For homotypic collectives, cells were trypsinized and seeded at the appropriate density on the micropatterns in their culture medium. For heterotypic doublets, after trypsinization, MEFs and HUVECs were mixed at a ratio of 1:2 using the HUVECs culture medium and seeded on the micropatterns. In both cases, cells not attached to the micropatterns were removed by washing with medium after 30 min and placed on the microscope for imaging.

### Cytoskeletal Drug Treatments

All the chemical inhibitors were added to the cells shortly after seeding (30mins) and kept in the medium during the entire time of observation. Concentrations of Rho Kinase Inhibitor (Calbiochem 555550) and Calyculin A (Sigma 208851) are indicated in the figures.

### Immunoflurescence

Cells were fixed for 10 to 15mins at room temperature in cytoskeleton buffer (CB = MES 10mM, KCl (ROTH Art. No. 6781.1) 138mM, MgCl (ROTH Art. No. KK36.1) 3mM, EGTA (Sigma E0396) 2mM in Milli-Q water) supplemented with Sucrose (ROTH Art. No. 4661.2) 10%, Triton-X100 (EUROMEDEX 3617818) 0.1%, Glutaraldehyde (Polysciences 00216) 0.1%, and Paraformaldehyde (Electron Microscopy Sciences 15710) 4%. When staining for p-MLC, fixation was preceded by a brief permeabilization step using Triton-X100 0.05% in CB supplemented with Glycerol (Carlo Erba 453742) 10%. The CS were rinsed 3 times with PBS. Aldehyde functions were then reduced using sodium borohydride (NaBH4 Sigma 452882) 1mg/ml in PBS for 10mins at room temperature and washed 3 times with PBS. The blocking solution (Bovine Serum Albumin (BSA A7030) 3% - Tween-20 (Sigma P2287) 0.1% - PBS) was then added for 45 min at room temperature. After that, the CS were successively incubated with the primary and secondary antibodies diluted in the blocking solution for 1hr each at room temperature. After both incubations, the CS were rinsed 3 times using PBS-Tween 0.1%. Then, the CS were labelled with a mixture containing Phalloidin Alexa Fluor 488 (Invitrogen A12379) at 1/200 and Hoechst (Invitrogen H3570) diluted in the blocking solution. Finally, the slides were rinsed 3 times in PBS-Tween 0.1%, once in PBS, and once in Milli-Q water and were then mounted in Mowiol 4-88 containing TetraSpeck™ Microspheres 0.5μm (Invitrogen T7281) at 1/200.

Rabbit Anti-Phospho-Myosin Light Chain 2 (Ser19) Antibody (Cell Signaling 3671) was used at 1/100 and secondary Goat anti-Rabbit IgG Alexa Fluor 546 antibody (Invitrogen A11035) was used at 1/500.

### Image Acquisition

Time-lapse imaging of the cells rotation was performed on an epifluorescence system consisting of a Nikon inverted microscope (Ti2) equipped with a Prime BSI Express Camera (Photometrics), a pE-4000 Fluo lamp (CoolLED), and a top stage incubator for controlled temperature 37OC and CO2 5% (Okolab). The following objectives were used: 4X Phase Contrast, 20X DIC, and 40X DIC. Time-lapse courses were followed using the NIS elements software.

Imaging of the cells rotation associated to traction force measurements was performed on a Nikon confocal spinning-disk system (Eclipse Ti-E) equipped with a CSUX1-A1 Yokogawa confocal head, an Evolve EMCCD camera (Photometrics), and a top stage incubator for controlled temperature 37OC and CO2 5% (TOKAI). The microscope was operated using MetaMorph software (MetaImaging). Phase Contrast 40X (air) or 40X oil objectives were used for the acquisition of images and movies.

Images of the immunofluorescence stainings were acquired on a Zeiss LSM900 Airyscan 2 confocal microscope (Axio Observer) using 63X objective (Plan-Apochromat 63X/1.4 oil).

In all the cases, only cells that were fully spread on the micropatterns were selected for imaging.

### TFM Analysis

Following the method previously described by (57), TFM data was analyzed in Fiji (58) using a homemade macro that takes advantage of a set of plugins (PIV and FTTC) (https://sites.google.com/site/qingzongtseng/imagejplugins). Displacement fields were obtained from fluorescent bead images before and after the detachment of the cells by trypsin treatment. Bead images were paired and realigned using template matching. Displacement fields were calculated by particle imaging velocimetry (PIV), which used a normalized cross correlation–based method with an iterative scheme. Final vector-grid size was 4.6μm × 4.6μm.

Only doublets fully spread on the micropatterns and rotating for at least 10 frames, corresponding to about 3hrs of rotation were selected for further analysis.

The TFM analysis was performed in two steps. First, Fourier-transform traction cytometry was used to compute the traction force field, with a regularization parameter of 5 × 10^−10^, Poisson coefficient of 0.5, and a Young Modulus of 16.7kPa. A circular region of interest was defined around the doublets, such that its area was larger than that of the micropattern (ROI ∼93μm in diameter). Force vectors located outside this ROI were discarded during the calculation of the doublets stored mechanical energy.

The second step consisted of extracting the magnitude of forces exerted by individual cells by reprocessing the previously generated traction force files and energy maps based on the manual definition of the cell-cell junction at every time point (59). Briefly, the defined force region was split at the junction into two areas, and then the corresponding forces were assessed. The sum of the TF vectors under each area represented the imbalance or the intercellular force. As the sum of the TF vectors in a doublet was zero (accounted for by the mechanical equilibrium of the doublet as a whole), the imbalance across both cells was equal in magnitude and opposite in direction.

In the process, the junction was fitted to an ellipse, whose long axis was rescaled to the size of the micropattern and used to compute the instantaneous angular displacement that provided both the direction and magnitude of doublet angular velocity.

For representing the average energy maps, the stored mechanical energy was projected overtime. The maps of distinct doublets rotating were then averaged.

For ME asymmetry visualization, the ROIs obtained by contouring the cells were used to realign the energy map at each time point, before projecting them overtime. This method was used for all the rotating doublets of each subpopulation, such that all the cells displaying the higher ME in average, termed “the stronger cell”, were located to the right (Movie 6). Finally, the resulting realigned energy maps of all CW- and CCW-rotating doublets were separately projected to generate the aligned, average energy map corresponding to each subpopulation.

### Image Analysis

Analyses were performed using Fiji software.

#### Quantification of Rotation and Bias of Doublets

Time-lapse movies, generated in Fiji ImageJ, were examined by the user, who manually identified the rotating collectives and indicated their directionality. To decrease the influence of inherent user bias on the outcomes, the analysis (blind or not) of the data generated from different experiments was performed by two, three, or four independent users.

Alternatively, for doublets, a macro was used to track the nuclei of the doublets and identify their rotational bias. Briefly, a threshold was first applied to the two detected nuclei and a binary mask was created. The nuclei were detected using particle analysis then tracked during the rotation of the doublets by minimal distance between two time points. The tracking process was semi-automatized to leave the possibility to refine the detection of the nuclei when less or more than two nuclei were detected.

Collectives, in which the cells retracted from the micropatterns or reverted their direction of rotation, were eliminated from the population that was subsequently used to quantify the statistics.

The percentage of rotating collectives within the entire cell population was computed. The bias was then quantified amongst the subpopulation of rotating collectives.

#### Generation of Vector Plots

For big collectives, tracks were computed using Trackmate plugin in Fiji. Parameters were fine-tuned for each tracking experiment to follow optimally each cell trajectory. Aberrant tracks (when nuclei detection was lost over time or upon inaccurate detection) were manually discarded. Nuclei trajectories were represented by color-coded line on defined periods specified in the figures. The representations display the end-point of the trajectories (circles). The start- and end-points of the tracks were subsequently used to generate vectorial displacement maps in Fiji using the plot function.

#### Actin and phosphomyosin Immunofluorescence Signal Quantification

Following inhibitors treatment and immunofixation, the total myosin signal was measured in Fiji on single plane images after contouring the doublets and then normalized by the intensity of tetraspeck fluorescent beads of 1 µm (Invitrogen T7282).

For immunofluorescence quantifications in HUVECs and MEFs homotypic doublets, a maximum projection of the acquired Z-stacks was first performed. The total actin and phosphomyosin signals were measured in a region of interest of constant area and then normalized by the intensity of tetraspeck fluorescent beads of 0.5 µm (Invitrogen T7281).

### Data representation and Statistical Analysis

Energy maps, traction stress maps, and vector plots were created in Fiji.

Data plotting, graph design, frequency distributions, and probability distributions were done on Graphpad Prism 8 (http://www.graphpad.com). The error bars represented on the plots correspond to standard deviations.

The statistical tests, indicated in each figure legends, were all performed on Graphpad Prism 8. For unpaired t-tests, a non-parametric Mann Whitney test was used.

## Data Availability

All data that support the findings of this study are available within the main text and SI of this paper.

## Acknowledgments

We thank Dr. Calina Copos for fruitful discussions and assistance on the statistical evaluation of our data. We also thank the microscopy facility MuLife of IRIG/DBSCI, funded by CEA Nanobio and GRAL LabEX (ANR-10-LABX-49-01) financed within the University Grenoble Alpes graduate school CBH-EUR-GS (ANR-17-EURE-0003). This work was supported by Agence Nationale de la Recherche (ANR-20-CE13-0004) and ARC (Association pour la Recherche contre le Cancer) foundation (ARCDOC42022120005746).

## Author Contributions

L.K, M.T. and L.B. designed research; G.B., L.K. and P.S. performed research; B.V., G.B., A.S., P.S. and L.K. analyzed data; G.B., L.K. and M.T. wrote the paper with input from all authors.

