## Supplementary figures and movies for "Contractile forces direct the chiral swirling of minimal cell collectives"

### Supporting Information for
### Contractile forces direct the chiral swirling of minimal cell collectives

Ghina Badih[a], Alexandre Schaeffer[b], Benoît Vianay[a], Pauline Smilovici[a], Laurent Blanchoin[a,b], Manuel Thery[a,b,*], Laëtitia Kurzawa[a,*]
a Université Grenoble-Alpes, CEA, CNRS, INRA, Interdisciplinary Research Institute of Grenoble, CytoMorpho Lab, Grenoble, France.;
b Université Paris Sciences et Lettres, CEA, ESPCI, Institut Pierre-Gilles de Gennes, CytoMorpho Lab, Paris, France.

*Manuel Thery, Laëtitia Kurzawa.

**This PDF file includes:**

Figures S1 to S5
Legends for Movies S1 to S9

**Fig. S1**

A. Schematic representation of the experimental setup used for assessing the rotation of collectives. HUVEC cells collectives of different sizes were seeded on disk shaped-micropatterns (250µm or 60µm) coated with fibronectin.
B. Top panel: Time series of HUVEC cell collectives rotating CW (highlighted in purple) or CCW (highlighted in orange). Nuclei are stained with Hoechst (magenta) and trajectories determined in TrackMate over the time frames indicated on the images are represented by tracks on the images (the circles around the nuclei indicate the last time point of the trajectories). Panel below: corresponding vector plots of the trajectories. Scale bar=30µm.
C. Number of nuclei per collectives determined on the first frame. N=1; n=64 collectives. Statistical significance was assessed using unpaired t-test ($p=0.9281$).
D. Top image: Representation of the nuclei within a rotating collective. The trajectories of the nuclei determined in TrackMate are overlaid onto the nuclei images. The color code of each track encodes for the mean speed measured over the rotation (14hrs). Image below: Maximal projection image of the nuclei of the collective overtime. The yellow circle defines the boundaries of the regions in which the mean speeds of nuclei were measured. (Image scale bar =50µm; mean speed scale bar=µm/sec).
E. Mean speeds of nuclei located in the inner region of the collectives (IN) and in its outer region (OUT). N=1; n=64 collectives. Statistical significance was assessed using unpaired t-test ($p \leq 0.0001$).
F. Time-lapse sequences of different doublets populations: non-rotating (top panels), CW-rotating (middle panels), and CCW-rotating (bottom panels). Nuclei are labelled with Hoechst (magenta). Each nucleus is marked by a different colored spot and is identified overtime in the sequence.
G. Percentage of rotating and non-rotating HUVEC doublets Statistical significance was assessed using unpaired t-test ($p \leq 0.0001$).
H. Percentage of CW and CCW-rotating doublets. N=16 independent experiments; n=2295 doublets. Statistical significance was assessed using unpaired t-test ($p \leq 0.0001$).
I. Percentage of rotating doublets at 15hrs and 39hrs. N=1 experiment; n=271 doublets. Statistical significance was assessed using unpaired t-test ($p=0.0011$).
J. Percentage of CW-rotating doublets at 15hrs and 39hrs. N=1 experiment; n=181 doubets. Statistical significance was assessed using unpaired t-test ($p>0.9999$).

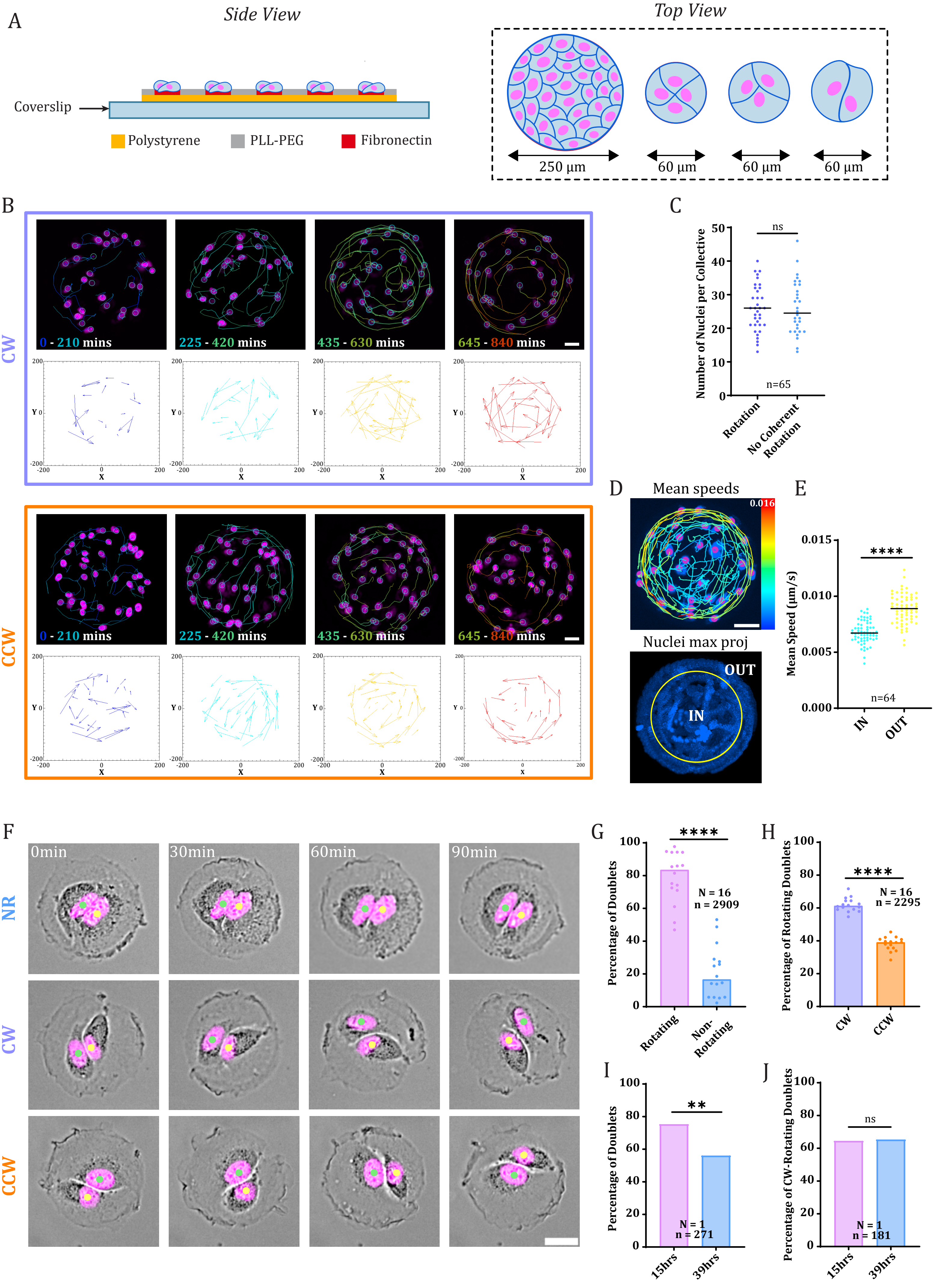
A
Side View
Top View
Coverslip
Polystyrene
PLL-PEG
Fibronectin
250 µm
60 µm
60 µm
60 µm
B
CW
CCW
0 - 210 mins
225 - 420 mins
435 - 630 mins
645 - 840 mins
C
Number of Nuclei per Collective
ns
n=65
Rotation
No Coherent Rotation
D
Mean speeds
0.016
Nuclei max proj
OUT
IN
E
Mean Speed (µm/s)
****
n=64
IN
OUT
F
0min
30min
60min
90min
NR
CW
CCW
G
Percentage of Doublets
****
N = 16
n = 2909
Rotating
Non-Rotating
H
Percentage of Rotating Doublets
****
N = 16
n = 2295
CW
CCW
I
Percentage of Doublets
**
N = 1
n = 271
15hrs
39hrs
J
Percentage of CW-Rotating Doublets
ns
N = 1
n = 181
15hrs
39hrs

**Fig. S2**

A. Time series of force stress maps overlaid onto phase contrast images of doublets rotating CW (above) and CCW (below). Traction force scale bar in Pa.
B. Representation of the corresponding Stored Energy maps overtime. To the right: representation of the time-averaged Stored Energy map. Color-coded scale bar in J.
C. Time series of the corresponding CW- (top images) and CCW-rotating doublets (bottom images) overlaid with the ROIs used to determine the mean junction length. On the right: plot of the mean junction length for CW- and CCW- rotating doublets. N=3 independent experiments; n=97 doublets. Statistical significance was assessed using an unpaired t-test (p=0.0061).

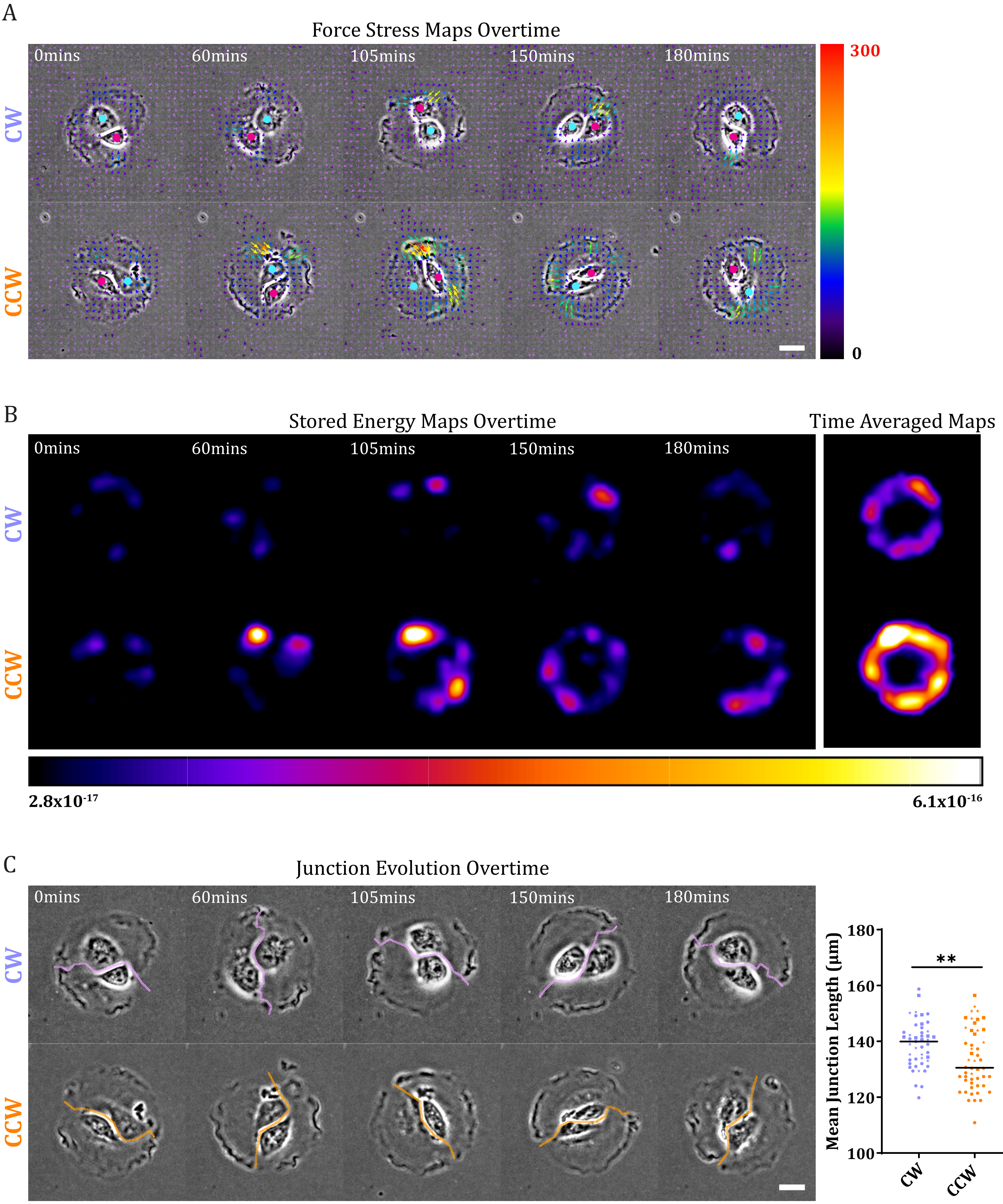
A
Force Stress Maps Overtime
0mins
60mins
105mins
150mins
180mins
CW
CCW
300
0
B
Stored Energy Maps Overtime
Time Averaged Maps
0mins
60mins
105mins
150mins
180mins
CW
CCW
2.8x10^-17
6.1x10^-16
C
Junction Evolution Overtime
0mins
60mins
105mins
150mins
180mins
CW
CCW
Mean Junction Length (µm)
180
160
140
120
100
**
CW
CCW

**Fig. S3**

A. Immunostaining of control (top images) and 20μM blebbistatin-treated (bottom images) HUVEC doublets. Stainings: F-actin and nuclei (left column); p-MLC (right column). Scale bar=10μm
B. Percentage of rotation quantified in control doublets and in doublets treated with increasing concentrations of Blebbistatin (10 and 20μM). N indicates the number of individual experiments and n the total number of doublets used for quantifications. Statistical significance was assessed using Chi-square test (Fischer's exact; Significance testing: ***=0.0001; ****<0.0001).
C. Percentage of CW-rotating doublets in control condition and with increasing concentrations of Blebbistatin (10 and 20μM). N indicates the number of individual experiments and n the total number of doublets used for quantifications. Statistical significance was assessed using Chi-square test (Fischer's exact; Significance testing: ns=0.0560; *=0.0148).

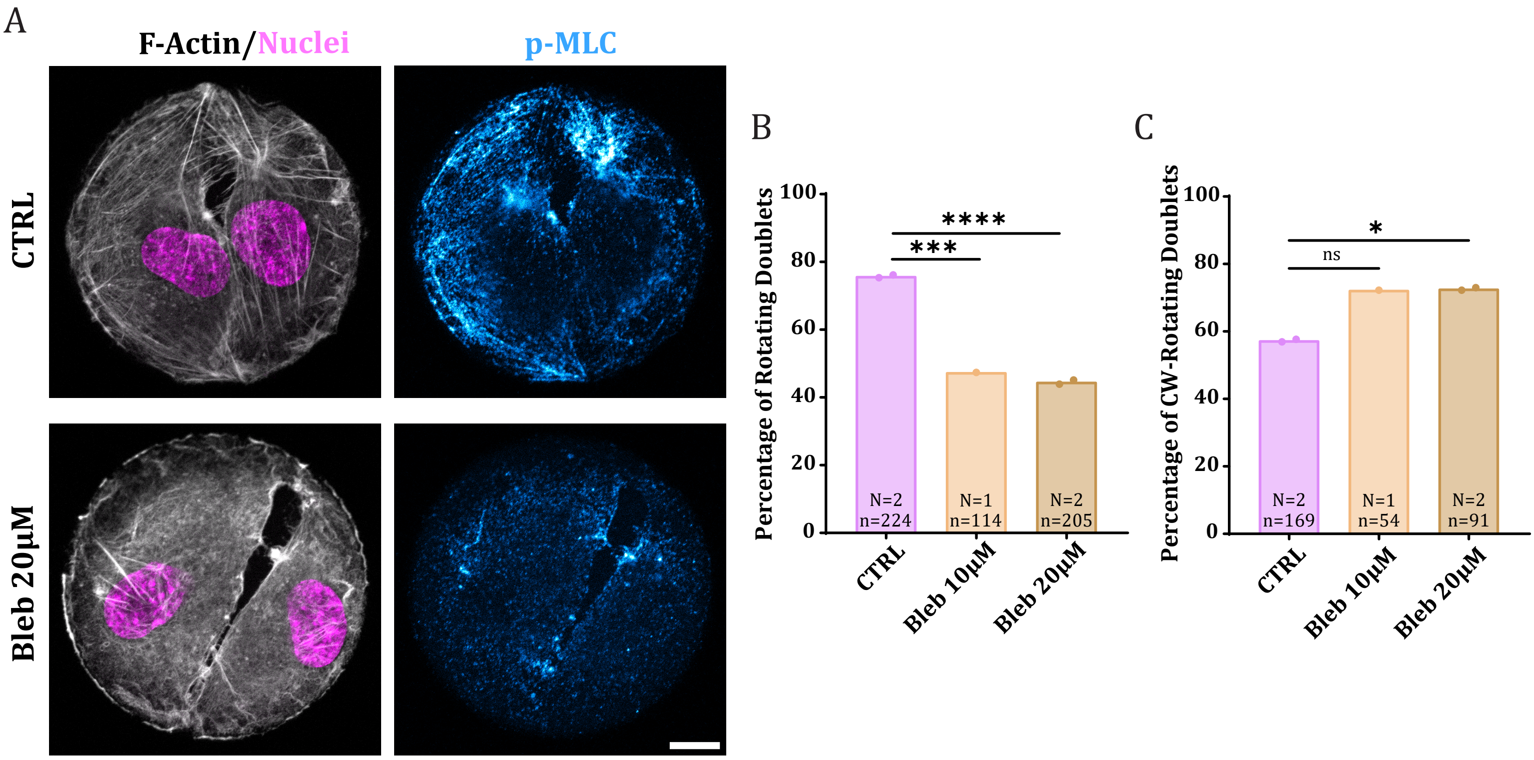
A
F-Actin/Nuclei
p-MLC
CTRL
Bleb 20µM
B
Percentage of Rotating Doublets
100
80
60
40
20
0
****
***
N=2
n=224
N=1
n=114
N=2
n=205
CTRL
Bleb 10µM
Bleb 20µM
C
Percentage of CW-Rotating Doublets
100
80
60
40
20
0
*
ns
N=2
n=169
N=1
n=54
N=2
n=91
CTRL
Bleb 10µM
Bleb 20µM

**Fig. S4**

A. Mean angular velocity of the rotating doublets as a function of the stored ME of their Weaker constituting cell. CW rotating doublets are represented in purple and CCW rotating doublets in orange. The Pearson correlation coefficient r is indicated on the plot. N=3; n98 doublets. A linear regression fit was applied on the data.
B. Relative frequency distribution of CW (purple) and CCW (orange) rotating doublets as a function of the stored ME of their Weaker constituting cell. Frequencies were calculated on $n_{CW}$=47 cells; $n_{CCW}$=50 cells.

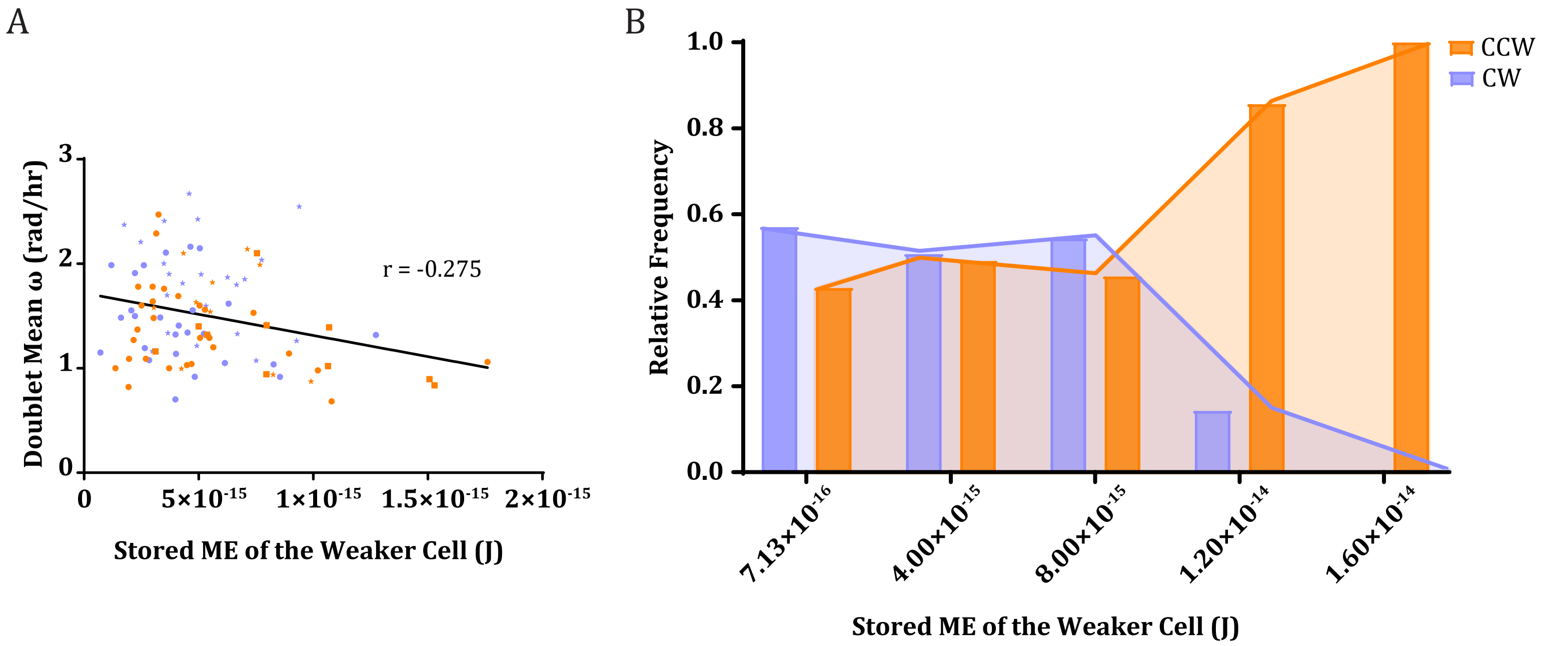
A
B
3
2
1
0
Doublet Mean ω (rad/hr)
r = -0.275
0
5×10-15
1×10-15
1.5×10-15
2×10-15
Stored ME of the Weaker Cell (J)
1.0
0.8
0.6
0.4
0.2
0.0
Relative Frequency
CCW
CW
7.13×10-16
4.00×10-15
8.00×10-15
1.20×10-14
1.60×10-14
Stored ME of the Weaker Cell (J)

**Fig. S5**

A. Images of distinct homotypic HUVEC and MEF doublets immunostained for F-actin (phalloidin in white), nuclei (Hoechst in purple), and phospho-myosin (p-MLC) in cyan.
B. Quantification of the F-Actin intensity signal for HUVEC and MEF doublets after normalization by the fluorescence of fiduciary markers. N=1; n=51 doublets for HUVEC and 50 for MEF. Statistical significance was assessed using an unpaired t-test ($p<0.0001$). Scale bar=10μm.
C. Quantification of the p-MLC intensity signal for HUVEC and MEF doublets after normalization by the fluorescence of fiduciary markers. N=1; n=51 doublets for HUVEC and 50 for MEF. Statistical significance was assessed using an unpaired t-test ($p<0.0001$). Scale bar=10μm.
D. Top images: Force stress maps of distinct homotypic MEF doublets of overlaid with phase contrast images at a single time point. Traction force scale bar in Pa. Bottom images: Corresponding Stored Energy maps. Color-coded scale bar in J.
E. Mean linear velocities measured in HUVEC and MEF homotypic rotating doublets. N=1. n= 284 doublets for HUVEC and 324 for MEF. Statistical significance was assessed using unpaired t-test ($p<0.0001$).
F. Representative images of distinct heterotypic HUVEC-MEF doublets immunostained for F-actin (phalloidin in white), nuclei (Hoechst in purple), and phosphor-myosin (p-MLC) in cyan (all signals are overlaid).

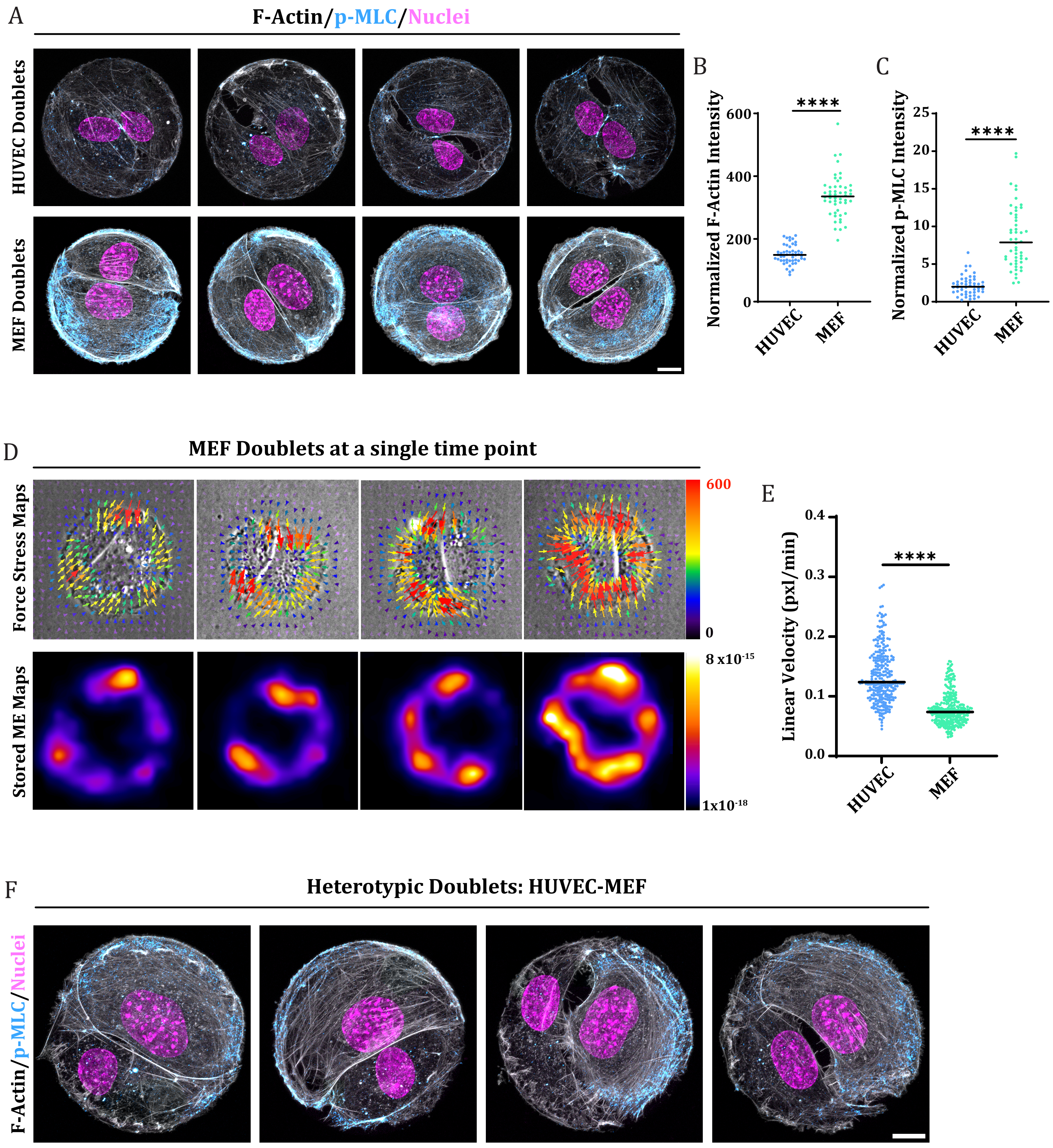
A
F-Actin/p-MLC/Nuclei
HUVEC Doublets
MEF Doublets
B
Normalized F-Actin Intensity
600
400
200
0
****
HUVEC
MEF
C
Normalized p-MLC Intensity
25
20
15
10
5
0
****
HUVEC
MEF
D
MEF Doublets at a single time point
Force Stress Maps
Stored ME Maps
600
0
8 x10-15
1x10-18
E
Linear Velocity (pxl/min)
0.4
0.3
0.2
0.1
0.0
****
HUVEC
MEF
F
Heterotypic Doublets: HUVEC-MEF
F-Actin/p-MLC/Nuclei

**Movie 1. Reorganization of HUVEC cells on disk of 250 µm diameter and onset of rotation.** Tracks determined in TrackMate are displayed on top of nuclei (Hoechst staining in white). Time frame: 1img/15min.

**Movie 2. Examples of CW- and CCW-rotating big collectives on 250 µm disks.** Overlays of HUVEC cells imaged in phase contrast and their nuclei (Hoechst staining in blue). Time frame: 1img/15min.

**Movie 3. Examples of CW-rotating doublet, triplet and quadruplet on 60 µm disks.** Overlays of HUVEC cells imaged in phase contrast and their nuclei (Hoechst staining in magenta). Time frame: 1img/15min.

**Movie 4. Examples of CW- and CCW-rotating doublets over 24h on 60 µm disks.** Overlays of HUVEC cells imaged in phase contrast and their nuclei (Hoechst staining in magenta). Time frame: 1img/15min.

**Movie 5. Examples of traction stress maps and stored energy maps obtained overtime for a CW- and a CCW-rotating doublets on 60 µm disks.** On the left: overlays of force stress vectors with cells imaged in phase contrast. Traction force scale bar=0-300 Pa. On the right: associated stored energy maps. Store Energy scale bar=$1.5e^{-17}$-$1.8e^{-16}$J. Time frame: 1img/15min.

**Movie 6. Examples of CW- and CCW-rotating doublets imaged in control conditions or in the presence of ROCKI at 7 µM.** Time frame: 1img/15min. Scale bar=30 µm.

**Movie 7. Examples of CW- and CCW-rotating doublets imaged in control conditions or in the presence of CalyA at 0.3 nM.** Time frame: 1img/15min. Scale bar=30 µm.

**Movie 8. Examples of traction stress maps before and after alignment of junctions and the corresponding aligned stored energy maps for a CW- and a CCW-rotating doublets on 60 µm disks.** On the left: overlays of non-aligned force stress vectors with cells imaged in phase contrast. Traction force scale bar=0-300 Pa. In the middle: overlays of aligned force stress vectors with cells imaged in phase contrast. On the right: associated stored energy maps. Store Energy scale bar=$5e^{-18}$-$5e^{-16}$J. Time frame: 1img/15min.

**Movie 9. Examples of CW- and CCW-rotating heterotypic doublets on 60 µm disks.** Overlays of cells imaged in phase contrast (+ green Calcein staining for MEFs only) and their nuclei (Hoechst staining in blue). Time frame: 1img/15min.
